# Inference of Diagnostic Markers and Therapeutic Targets from CSF proteomics for the Treatment of Hydrocephalus

**DOI:** 10.1101/2020.05.26.117457

**Authors:** Arie Horowitz, Pascale Saugier-Veber, Vianney Gilard

## Abstract

The purpose of this mini-review is to examine if publicly available cerebrospinal fluid (CSF) proteomics data sets can be exploited to provide insight into the etiology of hydrocephalus, into the character of the injury inflicted on the parenchyma by ventriculomegaly, and into the response of the brain to this condition. While this undertaking was instigated by reanalysis of recent comparative proteomics of CSF collected from the brain of healthy and *Mpdz* knockout (KO) mice (Yang et al., 2019), it is an opportunity to survey previously published CSF proteomics data sets to determine if they can be pooled together to that end. The overabundance of extracellular matrix (ECM) proteins, complement factors, and apolipoproteins in the CSF of *Mpdz* KO mice was taken to indicate that the hydrocephalic brain underwent ischemia, inflammation, and demyelination. The overabundance of five cytokine-binding proteins could be linked uniquely to insulin-like growth factor (IGF) secretion and signaling. The overabundance of two serpins, angiotensinogen and pigment epithelium-derived factor (PEDF) was considered as a biomarker of anti-angiogenic negative-feedback mechanisms to reduce CSF production. These findings raise the intriguing propositions that CSF proteomics can identify biomarkers of case-specific injuries, and that IGF signaling and angiogenesis pathways can serve as therapeutic targets. It appears, however, that the currently available proteomics data is not amenable to comparison of CSF from normal and hydrocephalic patients and cannot be used test the premise of those propositions.

## INTRODUCTION

Hydrocephalus is the most common cause of pediatric surgical intervention (Lim et al., 2018), occurring about once per 1000 births in the US (Kahle et al., 2016). Current treatments are invasive, involving either CSF drainage by ventriculo-peritoneal shunt (VPS), ventriculo-atrial shunt, and lumbo-peritoneal shunt, or endoscopic third ventriculostomy (ETV). Shunting entails increased rates of mortality and morbidity due to intraparenchymal hematoma during implantation and device failure or infection (Christian et al., 2016;Kahle et al., 2016;Riva-Cambrin et al., 2016). The hospitalization required for these procedures costs annually more than 2 billion dollars (Stein and Guo, 2008). CSF drainage from the ventricles to the peritoneum, the prevalent treatment for hydrocephalus, requires multiple revisions during the patients’ lifetimes at an average interval of 14.1 years (Reddy et al., 2014). Ultimately, surgical treatment is palliative and does not address the underlying cause of hydrocephalus. These exigencies provide the rationale for the implementation of non-invasive approaches in place of or in support of invasive treatments. To date, no effective non-invasive treatment has been reported.

Shunt implantation facilitates sampling and analysis of the CSF for early detection of infection (Khalil et al., 2016). It could conceivably be analyzed also for evaluating parenchymal health. If suitable markers are identified, their overabundance in the CSF of hydrocephalic patients could potentially serve as biomarkers of case-specific injuries. Furthermore, the composition of the CSF could provide information on the brain response to the elevated intra pressure exerted on the parenchyma as a result of ventriculomegaly. Delineation of the molecular pathways employed by the brain to this end could be used in principle to guide future pharmacological approaches and identify new druggable targets.

We had published comparative proteomics between the CSF of healthy and hydrocephalus-harboring *Mpdz* KO mice (Yang et al., 2019). The *Mpdz* knockout mouse model phenocopies similar loss-of-function mutations in the human ortholog, *MPDZ* (Al-Dosari et al., 2013;Saugier-Veber et al., 2017). Here, we compare the Yang et al. data on murine CSF to those of proteomic analyses of human CSF to gauge the similarity and, consequently, the validity of drawing diagnostic and therapeutic inferences collectively by pooling all the available data.

## ANALYSIS

### Inventory of relevant proteomic studies on human CSF

The number of comparative proteomic analyses of CSF of hydrocephalic patients to normal subjects reported to date is surprisingly small. We identified only three studies that had analyzed the CSF of hydrocephalus-harboring patients (Li et al., 2006;Scollato et al., 2010;Waybright et al., 2010). Out of these, only Li et al. compared the CSF of 15 adult patients with idiopathic normal pressure hydrocephalus (INPH) to 12 normal adult subjects. The CSF was resolved by 2D SDS-PAGE before undergoing mass-spectroscopy, reducing substantially the sensitivity of the assay and the total number of quantified proteins. Among the nine proteins they reported on, the intensity of the 2D ApoD spot was 2.3-fold higher in the CSF of INPH patients. The remaining eight proteins were not detected in our and others’ proteomic studies (see below). Scollato et al. also used 2D SDS-PAGE followed by mass-spectroscopy to analyze the CSF of 17 adult patients with NPH. They identified 12 proteins but did not compare the proteome of their CSF to that of normal subjects. Waybright et al. analyzed the CSF of nine infants, out of which one harbored congenital hydrocephalus, one underwent shunt revision, and the remaining seven acquired hydrocephalus caused by several types of CSF flow obstructions (Waybright et al., 2010). Unlike the previous studies, the CSF was processed by microcapillary liquid chromatography followed by mass spectrometry, resulting in higher sensitivity and culminating in the detection of 3336 proteins

Clearly, none of the previous studies affords comparison with the comparative proteomics we undertook (Yang et al., 2019). Similarly, we were unable to find published information on the specific functions of any of the 18 overabundant protein subgroup in the etiology of congenital, or, for that matter, any type of hydrocephalus. At most, the murine CSF proteome can be compared to the human one. To date, at least 31 studies undertook proteomics analysis of human CSF (see https://proteomics.uib.no/csf-pr/). We are aware only of the single aforementioned study by Waybright et al. for proteomic analysis of the CSF of hydrocephalic infants.

Based on their attributes, a subgroup of 18 proteins out of the 23 reported by Yang et al. was classified into five functional categories (from top to bottom in Table 1): extracellular matrix (ECM), complement factors, lipoproteins, cytokine binding proteins, and proteinase inhibitors. The 18 proteins were considered either as reporters of the injuries inflicted by ventriculomegaly on the parenchyma, or as components of autoregulatory pathways activated by the brain to counter these injuries. These classifications reflect literature searches on the associations between these proteins in the CSF of *Mpdz* KO mice (Yang et al., 2019) on one hand, and CNS pathologies that injured the parenchyma and affected CSF production, on the other. Five out of the original 23 overabundant proteins were excluded because we opted not to address the immune response, or because we did not find evidence in previous studies for their involvement in neuronal injury.

**Table 1:** List of proteins that were at least twice overabundant in the CSF of *Mpdz* KO mice relative to the CSF of healthy mice with a null hypothesis probability (*P*) of ≤ 0.05(Yang et al., 2019). The ratios of label-free quantification (LFQ) intensities provide the KO to wild type (WT) relative abundance of each protein (modified from Yang et al., 2019 (Yang et al., 2019)).

| Function | Name | Gene | LFQ KO/WT Ratio | *P*-value |
| --- | --- | --- | --- | --- |
| Extracellular matrix | Extracellular matrix protein 1 | Ecm1 | 11.691 | 0.0012 |
|  | Fibronectin | Fn1 | 10.589 | 0.015 |
|  | Gelsolin | Gsn | 4.415 | 0.0018 |
|  | Vitronectin | Vtn | 3.125 | 0.026 |
| Complement Factors | Complement C4-B | C4b | 5.121 | 0.0091 |
|  | Complement factor H | Cfh | 3.194 | 0.042 |
|  | Properdin | Cfp | 2.791 | 0.042 |
| | Fibrinogen $\beta$ chain | Fgb | 48.03 | 0.027 |
| Apolipoproteins | Apolipoprotein D | ApoD | 13.377 | 0.0003 |
|  | Apolipoprotein E | ApoE | 53.085 | 0.011 |
| Cytokine Binding | α-fetoprotein | Afp | 7.195 | 0.008 |
| | α-2-HS-glycoprotein | Ahsg | 5.664 | 0.024 |
|  | Insulin-like growth factor-binding protein 2 | Igfbp2 | 18.437 | 0.0005 |
|  | Insulin-like growth factor-binding protein 4 | Igfbp4 | 20.381 | 0.0003 |
|  | Sulfhydryl oxidase-1 | Qsox1 | 4.754 | 0.035 |
| Proteinase inhibitors | Angiotensinogen | Agt | 8.229 | 0.003 |
|  | Pigment epithelium-derived factor | Serpinf1 | 11.238 | 0.0009 |
| | α-2-macroglobulin | A2m | 37.142 | 0.0007 |

Out of the 18 overabundant proteins, 17 (except α-fetoprotein) were detected in the CSF of hydrocephalic patients (Waybright et al., 2010). Their majority (15) was among the top 100 (out of a total exceeding 3300) most abundant CSF proteins identified by Waybright et al. To further match the murine and human CSF proteomes, we searched the 18 proteins in the proteomic data sets from two studies on either 21 (Schutzer et al., 2010) or 38 (Lleo et al., 2019) CSF samples from normal adults. Either 16 (except fibronectin and α-fetoprotein) or 17 (except α-fetoprotein) proteins, respectively, were abundant in human CSF samples.

### Biomarkers of brain injury

The overabundant ECM proteins, complement factors and apolipoproteins (Table 1), were associated with mechanisms of ventriculomegaly-inflicted injuries based on the following evidence: ECM1 was linked to neuroinflammation, whereby it ameliorated the demyelination caused by T helper cells (Su et al., 2016). Plasma fibronectin, the soluble isoform detected in our previous study (Yang et al., 2019), supported neuronal survival and reduced brain injury following cerebral ischemia (Sakai et al., 2001). Extracellular gelsolin ameliorated inflammation (Bucki et al., 2008) and had a neuroprotective effect during stroke (Endres et al., 1999). Vitronectin activated microglia (Milner and Campbell, 2002), the primary immune effector cell of the CNS.

The complement system had a neuroprotective role in CNS neurodegenerative diseases (Bonifati and Kishore, 2007). Specifically, complement C4-B was produced by microglial cells in response to ischemia and inflammation (D’Ambrosio et al., 2001). Complement factor H was produced by neurons and had a protective function in neuroinflammation (Griffiths et al., 2009). Properdin (complement factor P), activated microglia and contributed to inflammation in response to ischemia (Sisa et al., 2019).

ApoE is produced by the choroid plexus and protects the brain against injury and promotes neuronal survival (Xu et al., 2006). ApoD has a similar neuroprotective role in CNS degeneration (Navarro et al., 2013). Lastly, α-2-macroglobulin was identified as a marker of neuronal injury (Varma et al., 2017).

Collectively, the overabundance of the aforementioned proteins reveals that severe hydrocephalus inflicted a set of injuries on the parenchyma, including neuroinflammation, demyelination, and impaired perfusion. Activation of CNS immune defense by microglial cells accompanied these processes and possibly aggravated the inflammation. This injury pattern conforms to the empirical picture of neonatal hydrocephalus (McAllister, 2012).

### Biomarkers of autoregulatory pathways

The IGF-binding proteins (IGFBPs) are produced in the CNS, including the choroid plexus (Tseng et al., 1989). Specifically, IGFPB2 and IGFBP4, which were overabundant in the CSF of *Mpdz* KO mice (Yang et al., 2019), are secreted by astrocytes in response to brain injury (Lewitt and Boyd, 2019). While the differential functions of each IGFPB are not fully understood, they are thought to stabilize IGF, extend its half-life (Zapf et al., 1986), and modulate its activity (Firth and Baxter, 2002). α-2-HS-glycoprotein, which binds to and modulates the signaling of the IGF receptor (Mathews et al., 2000), α-fetoprotein, a cytokine-binding plasma component (Mori et al., 2011), and sulfhydryl oxidase-1 participate in IGF uptake and transport by IGFPBs (Jassal et al., 2020). IGF and the IGFPBs are produced in choroid plexus epithelial cells (Stylianopoulou et al., 1988;Holm et al., 1994). The secretion of IGF into the CSF is increased in response to injury (Walter et al., 1999). Remarkably, all the five proteins in this group (Table 1) regulate multiple facets of IGF secretion and signaling. IGF protected neurons subjected to ischemia (Wang et al., 2013) and reduced microglia activation (Rodriguez-Perez et al., 2016).

Two of the overabundant proteins are the serpins angiotensinogen and PEDF (Table 1). Angiotensinogen is the precursor of several cleaved derivatives and peptides. One of the latter, angiotensin-2, reduced blood flow to the choroid plexus (Maktabi et al., 1990) and CSF production (Maktabi et al., 1993). However, its cleaved derivatives were antiangiogenic (Celerier et al., 2002). The anti-angiogenic is shared by PEDF (Dawson et al., 1999), which is secreted by ependymal cells in the subventricular zone (SVZ) (Ramirez-Castillejo et al., 2006). It appears, therefore, that serine protease inhibitors may reduce CSF production by pruning the dense capillary network that surrounds the choroid plexus vessels (Zagorska-Swiezy et al., 2008).

## DISCUSSION

Few previous studies had analyzed the diagnostic potential of the CSF. Developmental defects originating from attenuated cycling of germinal matrix cells in the brains of hydrocephalus-harboring rats had been attributed to alteration in the composition of the CSF relative to healthy subjects, but the putative difference between the CSF of health and hydrocephalic rats had not been identified (Owen-Lynch et al., 2003). Comparison of select proteins in the CSF of patients with idiopathic normal pressure hydrocephalus before VPS implantation and of healthy subject, was used to diagnose the damage to the patients’ parenchyma. Subsequently, the concentrations of the same proteins after VPS shunting were used to gauge if the shunt ameliorated the damage (Jeppsson et al., 2013). We did not find, however, published comparisons between the CSF proteomics of normal subjects and hydrocephalus-harboring patients that could be used for diagnostic or therapeutic purposes.

### Potential diagnostic implications

The interpretation of the overabundance of ECM proteins, complement factors, and apolipoproteins present in the subgroup of 18 proteins reported by Yang et al. as biomarkers of brain injuries is supported by previous studies (McAllister, 2012). Their individual or collective overabundances could be potentially measured in the CSF collected during VP shunting or ETV of hydrocephalic patients. Beyond verifying the occurrence of hydrocephalus, such measurements would highlight case-specific attributes, such as the presence of demyelination, revealed by the overabundance of ECM1, or of an aggressive immune response indicated by the overabundances of vitronectin and complement factors.

### Potential therapeutic implications

The combined overabundance of all the five cytokine-binding proteins present in the above subgroup suggests that IGF signaling is a major autoregulatory pathway employed by the brain, possibly to counteract the neuronal injury caused by ventriculomegaly (Johnston et al., 1996). Similarly, the concurrence of the overabundances of angiotensinogen and PEDF indicates that angiogenesis is suppressed as a negative feedback loop to counter the expansion of the ventricles by a reduction in CSF production. Each of these pathways could potentially be exploited for non-invasive therapies in place of or as adjuvants to the current invasive approaches. Finally, the marked overabundances of ApoE and ApoD suggest that these apolipoproteins may be used directly for neuroprotection of the parenchyma.

## List of abbreviations

Apo: apolipoprotein;
CNS: central nervous system;
ECM: extracellular matrix;
ETV: endoscopic third ventriculostomy;
ICP: intracranial pressure;
IGF: insulin-like growth factor;
IGFBP: IGF-binding protein;
KO: knockout;
LFQ: label-free quantification;
Mpdz: multi PDZ;
PDZ: PSD-95 (= postsynaptic density 95), Dlg (= discs large), ZO-1 (= zonula occludens-1);
PEDF: pigment epithelium-derived factor;
SVZ: subventricular zone;
VP: ventriculoperitoneal;
WT: wild type.

## Acknowledgement

The manuscript of this publication was released and posted on the bioRxiv preprint server (https://www.biorxiv.org/content/10.1101/2020.05.26.117457v3)

## Authors’ contributions

AH initiated the proteomics data analysis; all authors contributed to the writing of the manuscript.

## Ethics approval and consent to participate

N/A

## Consent for publication

All authors are in agreement.

## Availability of data and materials

N/A

